# Control of E-S Potentiation at two different sites in the dendro-somatic axis

**DOI:** 10.1101/2020.02.21.960138

**Authors:** Jason K. Clark, Daniel V. Madison

**Author notes:** Corresponding Author: DV Madison. The authors declare no competing financial interests.

## Abstract

EPSP-Spike (E-S) Potentiation occurs alongside synaptic Long-Term Potentiation (LTP), both triggered by high-frequency synaptic stimulation (HFS). In this study, we confirm the earlier findings that E-S potentiation appears to be prevented by prior reduction of GABA_A_ receptor-mediated inhibitory synaptic transmission. However, we demonstrate that this is a result of an occlusion of E-S potentiation, not a block. E-S potentiation and GABA_A_ antagonism each saturate postsynaptic action potential discharge, but E-S potentiation can still be induced by high frequency activation of synapses, even in the presence of pharmacological GABA_A_ blockade. These results suggest that GABA_A_ blockers/antagonists and E-S potentiation share an expression mechanism, namely the reduction of GABA_A_-mediated synaptic inhibition. We also assayed changes in the electrical coupling between dendrite and soma, and were surprised to find that this coupling is decreased following HFS, a change that would oppose E-S potentiation. This decrease in dendritic-soma electrical coupling (D-S coupling) was induced through the action of GABA_B_ receptors, but not maintained or expressed via the activity of these receptors. These data all together suggest that there are two distinct and opposing changes that occur as a result of HFS: 1) A decrease in passive dendro-somatic electrical coupling, and 2), an increase in coupling between the somatic EPSP and action potential generation. These two opposing influences may function as a homeostatic mechanism to balance the excitatory/inhibitory relationship between primary neurons and interneurons, and may represent a separate mechanism by which feedback and feed-forward synaptic inhibition can influence E-S coupling in opposite directions.

**Significance statement:** E-S Potentiation is an activity-dependent form of plasticity that boosts the efficiency of the coupling between synaptic input and action potential output in a neuron. Because it is induced by synaptic activity in series with the more familiar long-term potentiation (LTP), and is similarly persistent, it represents an additional mechanism by which memory traces may be stored within neural circuits. The significance of this paper is that it shows that there are at least two points of control for E-S potentiation which influence it in opposite directions, thereby providing additional basic mechanisms by which memory traces may be modulated.

## Introduction

High frequency activation of the Schaffer collaterals in Area CA1 of hippocampus, and other excitatory synapses in the hippocampal formation, results in the induction of at least two parallel types of synaptic potentiation: Long-Term Potentiation (LTP) of the Excitatory Post-Synaptic Potential (EPSP); and EPSP-Spike potentiation (E-S potentiation), the increase in postsynaptic action potential discharge for a given EPSP magnitude. That both LTP and E-S potentiation are induced simultaneously by the same high frequency activation of synapses was recognized in the earliest reports of LTP [1], although later reports have shown that it may be possible to induce them separately [2, 3].

Three broad mechanisms might account for the expression of E-S potentiation: 1) A decrease in GABA-mediated synaptic inhibition, which would act to decrease shunting of EPSP current at dendritic and/or somatic membrane on its way to the spike trigger zone at the axon hillock; 2) An increase in the electrotonic coupling between dendrite and soma; and 3) A change in the postsynaptic action potential threshold to more hyperpolarized voltages, which has been reported [4, 5], although this might be an indirect effect related to changes in synaptic inhibition. The other two mechanisms differ in whether they reflect a change in inhibitory neurotransmission, or are a cell-autonomous property of pyramidal neurons [6]. Previous studies have examined the role of dendritic electrical properties during production of E-S potentiation, but the conclusions have been inconsistent [7, 8].

E-S potentiation is generally thought to result at least in part from a reduction in GABAergic signaling. GABA_A_ antagonists appear to prevent the development of E-S potentiation [5, 6, 9–12]. Application of the GABA_A_ channel blocker Picrotoxin (PTX) before High Frequency Stimulation (HFS) potentiates the population spike signal, mimicking E-S potentiation, and this PTX enhancement is at least reduced by prior HFS [9]. Intracellular recordings show a decrease in the IPSP after tetanus that is blocked by application of the NMDA receptor antagonist AP5, and is Calcineurin dependent [12], but others show no significant change in IPSPs [13]. The induction of E-S potentiation, like LTP, seems to depend on the NMDA receptor, but it is not clear if these are the same populations of NMDA receptors for both plasticities. E-S potentiation may also be mediated by a long-term depression of synapses onto interneurons from pyramidal neurons [14]. In earlier studies, this was attributed to a decrease in feed-forward in inhibition, though feedback inhibition is more widely viewed as being mediated by GABA_A_ transmission, as opposed to a dominant role of GABA_B_ transmission in feed-forward inhibition [9, 15, 16]. The second major hypothesis for the development of E-S potentiation has involved potentially cell-autonomous mechanisms, such as changes in the intrinsic excitability or dendritic membrane resistance [4, 17–21].

In this study, we aim to explore the hypothesis that changes in inhibitory control contribute to the development of E-S potentiation and/or whether non-inhibition-related changes in electrotonic coupling might also participate. We find that blockade of GABA_A_ transmission occludes the expression of E-S potentiation, but not its induction. This occlusion occurs in both directions: prior application of a GABA_A_ blocker prevents HFS-induced increases in E-S coupling, and prior HFS prevents increases in coupling by application of a GABA_A_ blocker. But to our surprise, tetanic stimulation decreases the electrical coupling between dendrite and soma, a change that is in opposition to the production of E-S potentiation. This change is triggered by GABA_B_ transmission, and demonstrates a previously unreported form of GABA_B_ receptor-mediated intrinsic plasticity. These opposing effects may function as independent regulators of information processing or memory trace storage at the level of both the individual neuron, and the hippocampal micro-circuitry.

## Materials and Methods

### Animals and euthanasia

All animals used in this study were male Sprague-Dawley rats approximately 6 weeks old, obtained from Charles River Laboratories (Wilmington, MA). Animals were housed at Stanford University in an AAALAC accredited facility on a 12 hour light/dark timed scheduled with ad libitum access to food and water. Euthanasia of animals occurred under deep anesthesia with isoflurane followed by decapitation. All procedures were performed in strict compliance with a protocol approved by The Stanford University Animal Care and Use Committee.

### Chemicals and reagents

Except where noted, specialty chemicals were obtained from Sigma Aldrich (St. Louis, MO) or Tocris Biosciences (Minneapolis, MN).

### Extracellular field potential recording

Hippocampal slices were prepared from male wild-type Sprague-Dawley rats approximately 6 weeks old. Rats were deeply anesthetized with isoflurane prior to decapitation. The brain was removed and submerged in ice-cold, oxygenated (95% O_2_ / 5% CO_2_) dissection artificial cerebrospinal fluid (dACSF) containing: NaCl (119 mM), KCl (2.5 mM), MgCl_2_ (3.8 mM), NaH_2_PO_4_ (1.0 mM), NaHCO_3_ (26.2 mM), and D-Glucose (11 mM). The brain was sectioned using a Vibratome through the horizontal plane into 500 µm thick slices. The hippocampus was then dissected free from slices and the CA3 region was removed. We estimate such slices were from the middle 30% of the hippocampus with respect to the longitudinal axis. We excluded slices from the extreme dorsal and ventral poles. Slices were placed in a submersion recording chamber (Harvard Apparatus, Holliston, MA) and perfused at approximately 100 ml/hr (1.67 ml/min, Gilson miniplus 2 peristaltic pump) with oxygenated (95% O_2_ / 5% CO_2_) standard ACSF containing: NaCl (119 mM), KCl (2.5 mM), MgCl_2_ (1.3 mM), CaCl_2_ (2.5 mM), NaH_2_PO_4_ (1.0 mM), NaHCO_3_ (26.2 mM), and D-Glucose (11 mM). Slices recovered for 30 minutes at room temperature and an additional 120 minutes at 30°C.

A bipolar stimulating electrode (FHC, Inc., Bowdoin, ME) was placed within the stratum radiatum of CA1, and two glass capillary extracellular recording microelectrodes filled with 3M NaCl (pulled on a Sutter Instruments P-87 Brown flaming microelectrode puller to a resistance of approximately 5 MΩ) were placed within the hippocampal slice with one in the stratum radiatum layer of CA1, and the other in the stratum pyramidal layer of CA1. Both recording electrodes were placed in the same horizontal plane with each electrode tip aligned to the same imaginary longitudinal y-axis. Synaptic field Excitatory Post-Synaptic Potentials (fEPSPs) and simultaneously recorded somatic field population spikes were recorded in response to stimulation of Schaffer collateral→CA1 synapses using a stimulus pulse consisting of a single square wave of 100 µs duration. Data were collected and digitized at 10 kHz, low-pass filtered at 1 kHz, and analyzed with pCLAMP 11.0 (Molecular Devices, San Jose, CA) or by lab-written software under LabVIEW 7.1 (National Instruments, Austin, TX). The initial slope of the synaptic fEPSP was measured by fitting a straight line to a 1 ms window to the initial slope of the fEPSP. The slope of the somatic fEPSP was measured by fitting a straight line to a 1 ms window immediately prior to the onset of the population spike. The area of the population spike was measured by creating an imaginary line from the spike onset to the spike termination and capturing the area within the spike under the imaginary border (see Fig 1A). Area was measured instead of amplitude as an index of total neuron firing without regard to differences in synchronicity.

**Figure 1:**
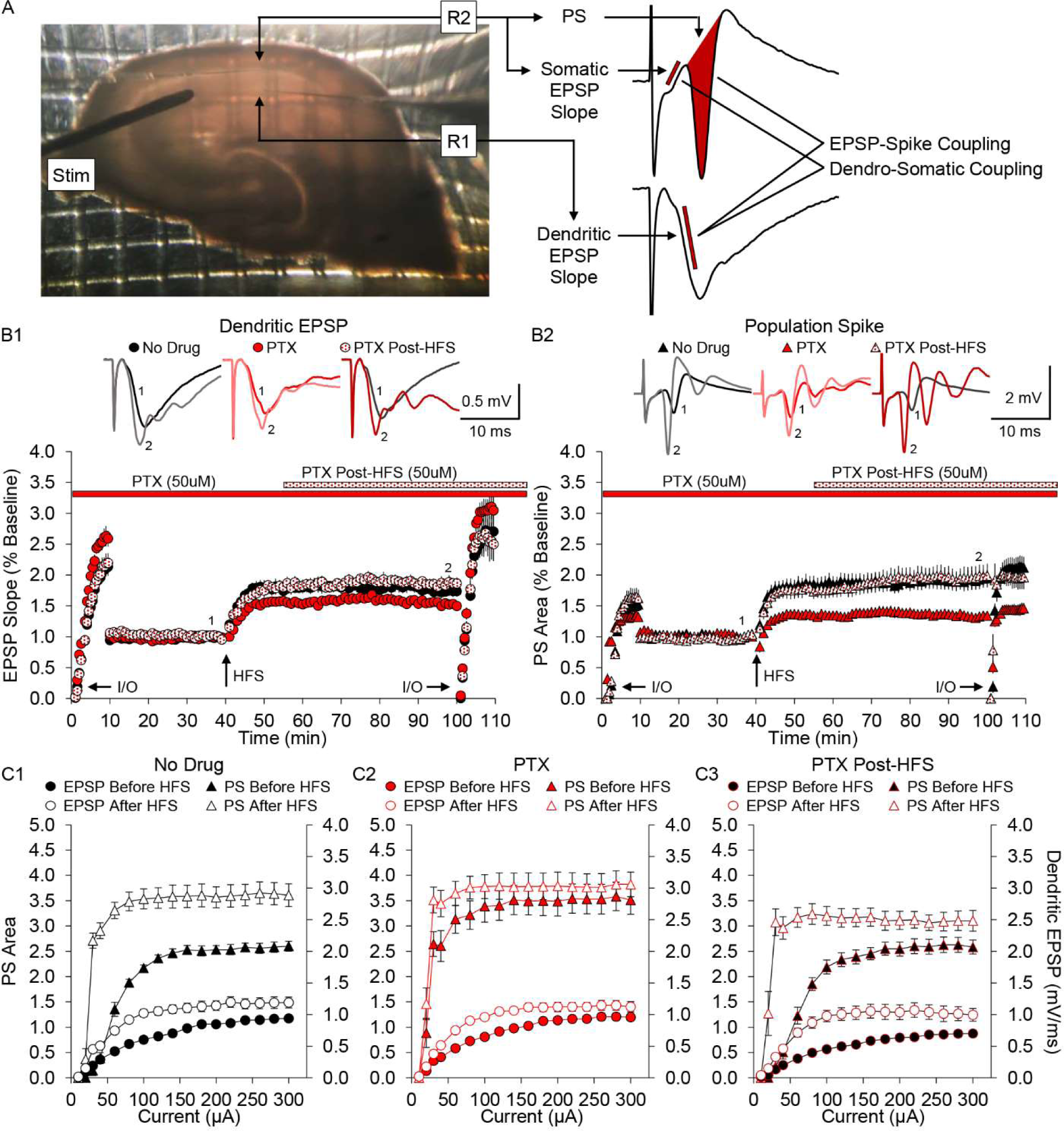
Picrotoxin reduces potentiation of the population spike when applied before, but not after HFS. A. A Hippocampal slice showing the placement of the stimulating electrode (Stim) in stratum radiatum near the CA2/CA1 border, and two recording electrodes (R1 and R2) in area CA1: one in stratum (s.) radiatum where Schaffer collateral synapses terminate on CA1 pyramidal cell dendrites (R1), and the other in s. pyramidal where the CA1 pyramidal cell bodies are located (R2). To the right are exemplar field potentials from these two electrodes showing the zone of measurement of the rising slope of the dendritic EPSP field (lower trace), the zone of measurement of the rising slope of the somatic EPSP field and the measurement of the area of the first population spike (upper trace). B1. The time-course of experiments showing the effects of HFS (up arrow) on the dendritic EPSP field in the presence of the GABA_A_ channel blocker Picrotoxin (PTX) (50 µM) applied either before or after HFS. Filled black symbols: No Drug control; red filled symbols: PTX present throughout experiment; red hatched symbols: PTX applied post-HFS. Time of delivery of PTX for each condition is indicated by the red bar (PTX present throughout experiment) or by the hatched bar (PTX applied 15 min post-HFS) located above the plot. An Input/Output curve was constructed at the beginning and end of each experiment (indicated by I/O arrows at the bottom of the plot). Application of PTX post-HFS resulted in a field EPSP that was higher than when PTX was applied throughout the experiment, though neither PTX nor PTX post-HFS groups were different than No Drug control (No Drug: 73 ± 8%, n=12; PTX: 52 ± 3%, n=12; PTX post-HFS: 86 ± 9%, n=10) as determined by ANOVA (F(2,31)=5.65, p < 0.01). B2. The population spike recorded simultaneously with the EPSPs in B1. PTX applied before HFS reduced the potentiation of the population spike by HFS, but had no effect when applied after HFS, which was similar to No Drug control (No Drug: 93 ± 15%; PTX: 33 ± 5%; PTX post-HFS: 97 ± 19%)(F(2,31)=7.38, p < 0.01). Averaged traces before (1) and after (2) HFS are displayed above the plots for each of the three conditions, and the corresponding time points for these averaged traces are indicated on the LTP time-course below. C. Input/Output curves constructed from a series of standardized stimulus strengths at the beginning and end of each experiment (visible in the LTP plots of panel B indicated by the I/O arrows). The two upper series in each graph are the population spike area (left axis on each graph), and the two lower series are the field EPSP slope (right axis on each graph). Each graph shows I/O curves before (filled symbols) and after (open symbols) HFS. Note: Recordings for each I/O curve in panel C are the same as in panel B. The I/O curves represented in the LTP time-course plots (panel B) are normalized values relative to baseline measurements before HFS, while values in panel C are the absolute values for population spike area and EPSP slope. Stimulus strengths are 10, 20, 30, 40, 60, 80, 100, 120, 140, 160, 180, 200, 220, 240, 260, 280, and 300 µA. Data shown represent the mean ± SEM.

Stimulus response (Input/Output) curves were obtained at the beginning and end of each experiment, and at specific time points within an experiment when appropriate as indicated for each figure. Stimulus pulses were delivered at 10, 20, 30, 40, 60, 80, 100, 120, 140, 160, 180, 200, 220, 240, 260, 280, and 300 µA at 0.033 Hz (once every 30 s). Initial conditions were set such that the population spike to dendritic fEPSP ratio was 3:1, to maintain dynamic range for the population spike, particularly below saturation. Long-Term Potentiation (LTP) experiments were carried out with a baseline stimulus delivered at 0.0167 Hz (once every 60 s) and normalized by dividing all synaptic fEPSP slope values or population spike area values by the average of the 5 responses recorded during the 5 minutes immediately prior to High Frequency Stimulation (HFS). LTP was induced by subjecting the slice to HFS consisting of 3 episodes of 100 Hz for 1 s stimulus trains (100 pulses x3) administered at 20 s inter-train intervals. Values indicating drug response were normalized to the 5 minutes of responses immediately prior to drug application, and were determined by averaging 5 minutes of normalized values at specific time points after drug application as indicated for each figure.

Due to the dynamic nature of E-S potentiation, quantification was estimated by determining the difference in population spike area before and after HFS with input matched dendritic EPSPs (i.e., we compared population spikes with an fEPSP slope of ∼0.7 mV/ms before HFS, to population spikes with an fEPSP slope of ∼0.7 mV/ms after HFS). Data used for this analysis was obtained from Input/Output curve data.

In order to evaluate the contribution of GABAergic signaling on LTP and E-S potentiation, the GABA_A_ channel blocker Picrotoxin (50 µM) or the GABA_B_ antagonist CGP-54626 (3 µM) (Tocris Bioscience, Minneapolis, MN) was bath applied 80 min prior to HFS. In experiments where Picrotoxin or CGP-54626 was applied after LTP induction, it was bath applied 15 min post-HFS as not to interfere with any post-tetanic potentiation mechanisms. In experiments where Picrotoxin was washed out after LTP induction, washout began 5 minutes post-HFS.

### Experimental Design and Statistical Analysis

Sample size was calculated with an expectation of being able to detect changes of 20% or more in measurements. Given the typical standard deviation of similar experiments, a power of 0.9, and an alpha cutoff of 0.05, this would require a sample size of approximately 8. Tests of significance were performed using a One-way ANOVA with Tukey’s post-hoc analysis for comparison of HFS induced potentiation of the dendritic EPSP slope, somatic EPSP slope, and population spike area across treatment groups. Analysis of the effect of drug application on recordings within a specific treatment group was performed using a paired t-Test. Data pertaining to the dendritic EPSP slope, somatic EPSP slope, and population spike area generated from input matched traces before and after HFS, and used to construct bar graph data, were compared using independent t-Test for each treatment group. Statistical analysis was performed with SYSTAT 13 software (San Jose, CA). Reported n-values indicate the number of slices and animals assessed (one slice/animal).

## Results

Electrophysiological recordings of extracellular field potentials were obtained by stimulation of the Schaffer collateral axons and recording of both the dendritic EPSP field in the s. radiatum and the resulting simultaneously recorded population spike field in the s. pyramidal layer, in the CA1 region of mid hippocampus (Figure 1A). In each experiment, LTP and E-S potentiation were induced simultaneously with the same High Frequency Stimulation (HFS: 100 Hz for 1 sec tetanic stimulation x3). We performed experiments designed to test whether the GABA_A_ open channel blocker Picrotoxin (PTX) prevented the induction of E-S potentiation by applying HFS in the presence or absence of PTX (50 µM) under the following three conditions: 1) No Drug (control, no PTX), 2) PTX bath-applied throughout the experiment (PTX), and 3) PTX applied beginning 15 min after HFS (PTX post-HFS) (Figure 1B). In every experiment we conducted an Input/Output (I/O) curve at the beginning and end of each experiment to determine the relationship between the EPSP and action potential generation (population spike) before and after the induction of potentiation by HFS (Figure 1B and 1C). HFS induced potentiation of both the dendritic EPSP (Figure 1.B1) and the population spike (Figure 1.B2) in all three conditions. Results showed LTP of the dendritic EPSP in both the PTX (n=12) and PTX post-HFS (n=10) groups was not different than the No Drug control (n=12). LTP of the population spike however, was reduced when PTX was applied throughout the experiment compared to control, but had no effect on the population spike when applied after HFS, which was similar to control (No Drug: 93 ± 15%; PTX: 33 ± 5%; PTX post-HFS: 97 ± 19%)(ANOVA, F(2,31)=7.38, p < 0.01). As expected, the I/O curves in all three conditions for both the dendritic EPSP field and the population spike were elevated at the end of the experiment after HFS, compared to the I/O curves at the beginning of the experiment before HFS (Figure 1C). Note that the I/O curves of both the EPSP field and population spike in appear on the same graph.

Data from the I/O curves in Figure 1C was used to construct E-S coupling curves for each of the three conditions. These curves were constructed by plotting the slope of the dendritic EPSP against the area of the population spike to show the relationship between synaptic input and spike output throughout a range of synaptic responses (Figure 2). In the absence of drug, HFS results in a persistent leftward shift in the E-S coupling curve, indicative that E-S potentiation has occurred (Figure 2.A1). When PTX was present throughout the experiment however, HFS produced no shift in the E-S coupling curve, indicating no change in E-S coupling and no E-S potentiation in this condition (Figure 2.B1), consistent with previously published results that blockade of GABA_A_ transmission blocks E-S potentiation [9]. However, we found that if PTX was applied after the induction of potentiation (post-HFS), that this produced only a minimal enhancement of that shift (Figure 2.C1). These results suggest one of two possibilities: 1) That PTX present during induction blocks the induction of E-S potentiation, or 2) PTX causes a change in the relationship between the EPSP and population spike that occludes the expression of E-S potentiation.

**Figure 2:**
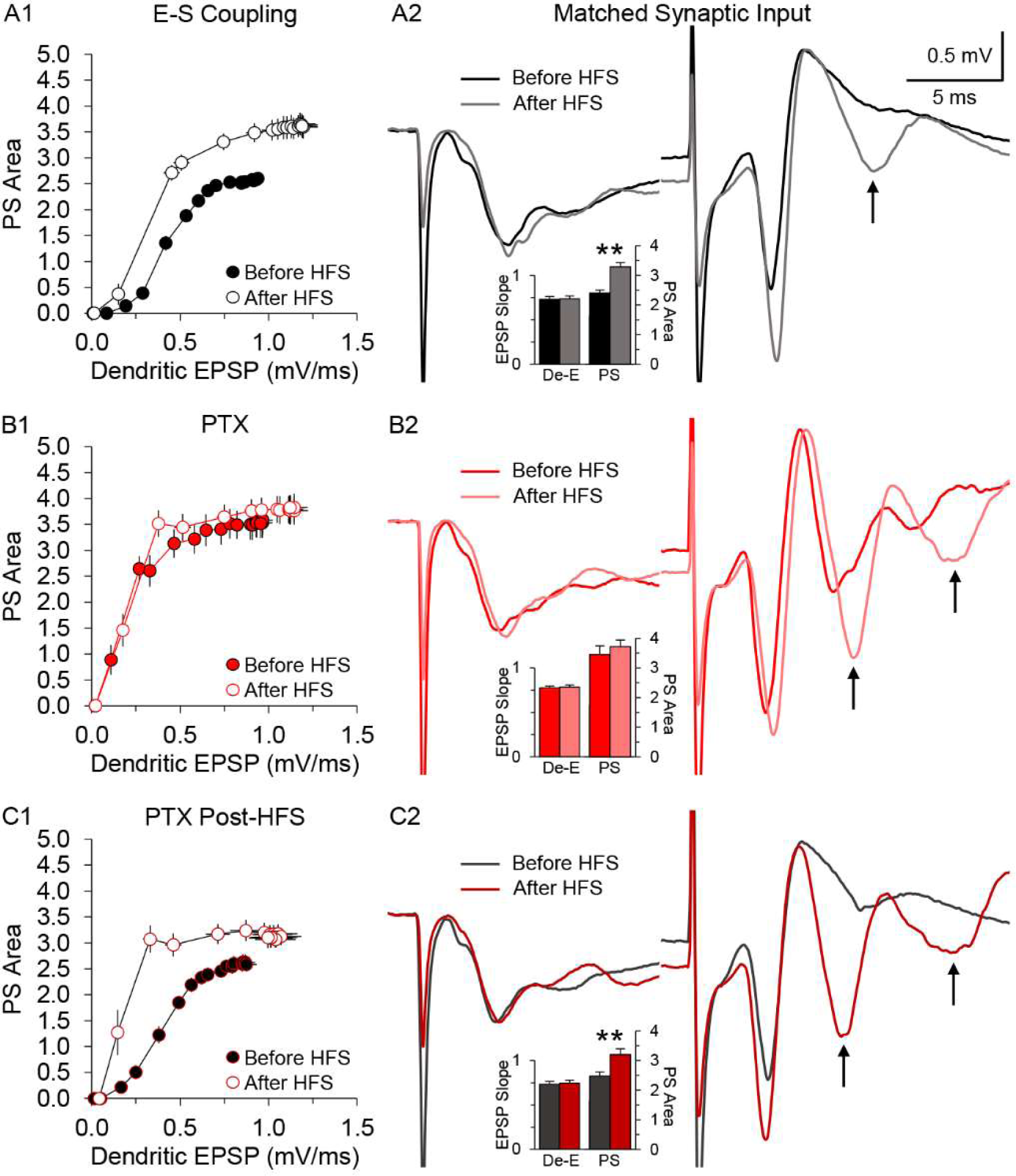
Shift in E-S coupling by HFS and Picrotoxin. The left column panels are E-S coupling curves constructed by plotting the population spike area against dendritic EPSP slope (see Figure 1A for reference), data taken from the I/O curves of figure 1C. A1. E-S coupling curves before and after HFS under No Drug control conditions (no PTX present). HFS causes a persistent leftward shift in the curve. A2. Averaged traces of input matched dendritic EPSPs (left traces) and the resulting simultaneously-recorded population spikes (right traces) before and after HFS. Traces after HFS have a reduced stimulus strength to match the rising slope of the dendritic EPSP to that of the dendritic EPSP before HFS. Dendritic EPSPs after HFS generate a larger population spike compared to matched dendritic EPSPs before HFS. This results in the persistent leftward shift in the E-S coupling curve throughout the range of synaptic input as shown in A1. The observed change in E-S coupling is indicative of EPSP-Spike potentiation, and is demonstrated by the larger population spike shown in A2. The inset bar graph in A2 shows the significant potentiation of the population spike (PS) with matched dendritic EPSPs (De-E) before (black bar) and after (dark grey bar) HFS (t-Test, p < 0.01). B1. E-S coupling curves with PTX present at the time of HFS and throughout the experiment. There is no leftward shift in the E-S coupling curve after HFS. B2. The population spike is not different with input matched dendritic EPSPs before and after HFS in the presence of PTX. There is no leftward shift in the E-S coupling curve throughout the range of synaptic input as shown in B1, indicating EPSP-Spike potentiation does not occur when PTX is present during HFS, also shown by the inset bar graph comparing responses before (red bar) and after (light red bar) HFS (t-Test, p = 0.48). C1. E-S coupling curves without PTX during HFS, but added post-HFS. HFS causes a persistent leftward shift in the curve, similar to that seen in control. C2. Input matched dendritic EPSPs produce a greater population spike after HFS compared to before. This is observed by the persistent leftward shift in the E-S coupling curve throughout the range of synaptic input, indicating EPSP-Spike potentiation occurs when PTX is not present during HFS, also shown by the inset bar graph comparing responses before (dark grey bar) and after (dark red bar) HFS (t-Test, p < 0.01). Note the afterdischarge spike(s), indicated by the arrows, in the population spike trace after HFS in all groups, thought to occur as a result of reduced inhibition following HFS (No Drug), or a combination of HFS and drug application (PTX, PTX post-HFS). Data shown represent the mean ± SEM. Bar graph data are generated from matched dendritic EPSP slopes and resulting population spike areas taken from Input/Output curves before and after HFS from panel 1C. Recorded traces for each matched pair were averaged and shown here in A2, B2, and C2 respectively for each group.

While LTP can be observed directly as the increase in both the dendritic EPSP and population spike, E-S potentiation is most clearly seen in the leftward shift of the E-S coupling curves, where it is clear that a given sized EPSP produces a greater population spike after HFS. E-S potentiation can also be observed in a second manner, by turning down the stimulus strength post HFS, to produce a dendritic EPSP field that matches the magnitude of the pre-HFS dendritic field. These input matched traces appear in Figures 2.A2, B2 and C2. The inset bar graphs show averaged values for EPSP slope and population spike area corresponding to the averaged traces shown above. Despite the reduced stimulus strength, matched dendritic EPSPs after HFS generate greater population spikes compared to matched dendritic EPSPs before HFS in both the No Drug control (Figure 2.A2; t-Test, p < 0.01), and when PTX is applied post-HFS (Figure 2.C2; t-Test, p < 0.01), showing that a given sized EPSP produces a greater spike output from that population of pyramidal neurons; i.e., E-S potentiation occurred. This is not seen however, when PTX is present throughout the experiment (Figure 2.B2; t-Test, p = 0.48), consistent with the E-S coupling curve in Figure 2.B1.

Upon closer inspection, we observed that the presence of PTX prior to HFS seemed to produce a population spike that was similar in magnitude to the population spikes after induction seen in the No Drug control and PTX post-HFS groups. In addition, the E-S coupling curve prior to HFS seen when PTX was present throughout the experiment (Figure 2.B1) was more similar to the leftward-shifted curves caused by HFS in the No Drug control and PTX post-HFS groups, than to the baseline non-shifted curves in those groups. This suggested that the effect of PTX may not actually prevent E-S potentiation, but rather occlude it, by pre-saturating the population spike through an increase in firing secondary to a reduction of GABA_A_-mediated synaptic transmission.

To differentiate between block or occlusion of E-S potentiation by PTX, we performed experiments to determine whether E-S potentiation could still be induced by HFS in the presence of PTX, even if its expression was occluded. We carried out this test by applying PTX to baseline recordings, then either applying HFS for induction, or not, before subsequently washing PTX from the slice. If the population spike remained elevated after washout of PTX, then E-S potentiation was induced, but if the population spike returned to baseline, then it was not (Figure 3). We conducted I/O curves at three different time points throughout the experiment: At the beginning, after the application of PTX but before the induction of potentiation with HFS, and at the end of each experiment after PTX washout. E-S coupling curves were constructed from these data in the same manner as in Figure 2.

**Figure 3:**
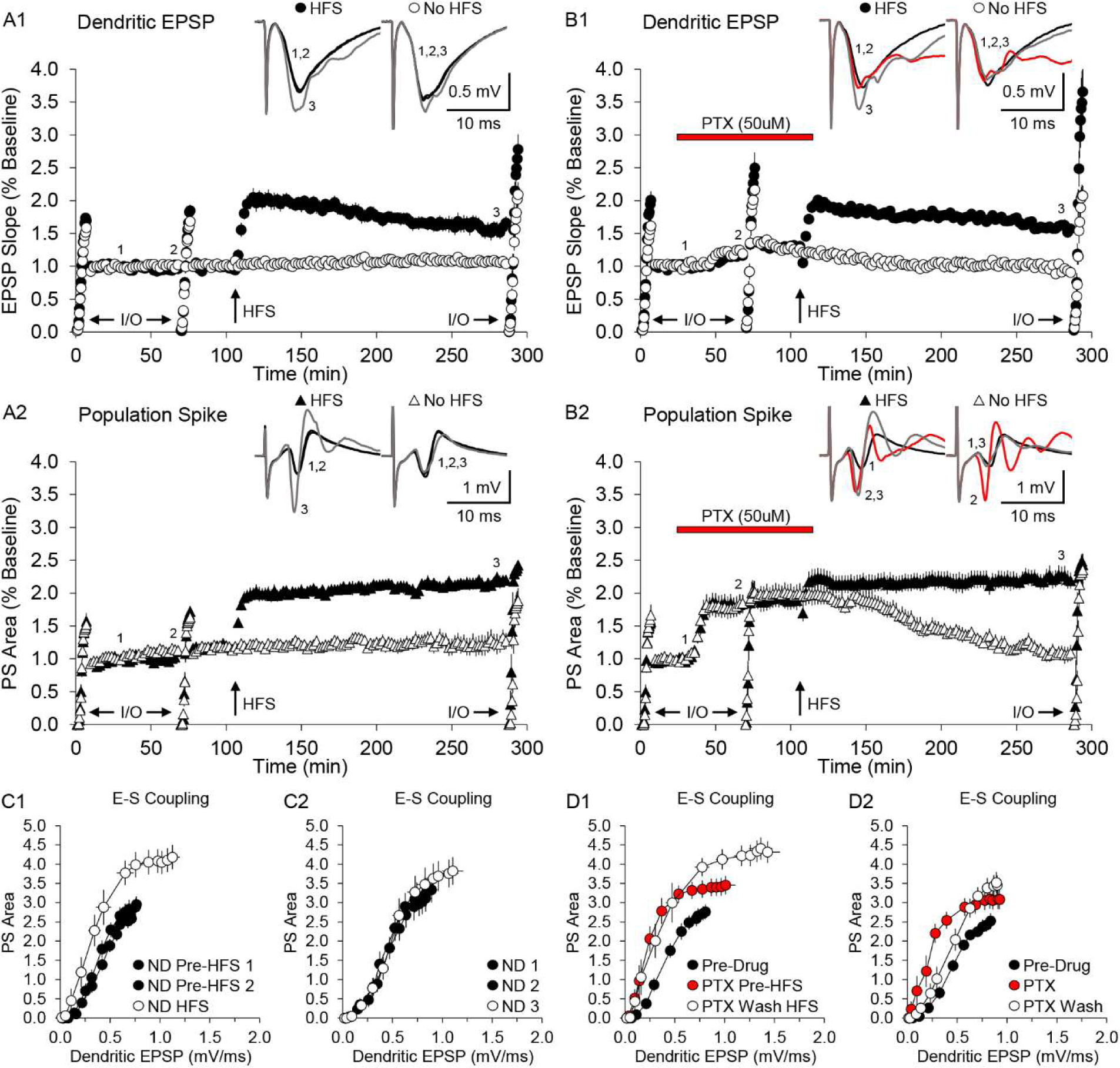
EPSP-Spike Potentiation is occluded, not blocked, by Picrotoxin. A and B. The effects of PTX application on basal activity of dendritic EPSPs and resulting population spikes, and subsequent washout either with or without HFS. Each experiment shows an HFS (filled symbols) vs. No HFS (open symbols) comparison. Three different Input/Output curves were constructed (indicated by the I/O arrows): at the beginning of the experiment, after the application of PTX (50 µM, red bar located above the plot) but before HFS, and at the end of each experiment. A. No Drug control condition for the dendritic EPSP (A1) and population spike (A2). B. Same experiment as in panel A, except with the application of HFS during application of PTX, then subsequently washed out (PTX-WO) starting five minutes post-HFS. Data shown in all experiments are normalized to the 5 minutes immediately prior to PTX application. The presence of PTX had no effect on the induction of LTP of the dendritic EPSP field (A1 vs B1), and following PTX washout, was not different from control (HFS - No Drug: 59 ± 10%, n=9; PTX-WO: 56 ± 7%, n=10). Experiments in which no HFS was performed showed responses after PTX washout were not different from control, indicating no lasting effects of PTX application only (no HFS – No Drug: 6 ± 6%, n=8; PTX-WO: −6 ± 6%, n=9). However, PTX caused a significant increase in the population spike (B2) compared to the same time point in the control (A2) (No Drug: 8 ± 6%; PTX: 81 ± 14%; t-Test, p < 0.01). HFS caused no further significant increase in the population spike, and upon PTX washout, remained potentiated, and was not different than the population spike after HFS in control (HFS – No Drug: 121 ± 7%; PTX-WO: 122 ± 13%). Without HFS, washout of PTX returned the population of spike to baseline and was not different from control conditions without HFS (no HFS – No Drug: 25 ± 13%; PTX-WO: 6 ± 12%). Averaged traces at three different time points are displayed above the plots for each condition, and the corresponding time points for these averaged traces are indicated on the LTP time-course below. The red trace in panel B indicates the presence of PTX. C and D. E-S coupling curves constructed from the experiments in panels A and B above. C1. No Drug control with HFS: 1^st^ and 2^nd^ E-S curves before HFS are similar, but following HFS there is a persistent leftward shift in the E-S coupling curve. C2. No Drug control without HFS: There is no change in E-S coupling throughout the duration of the experiment. D1. PTX with HFS: Application of PTX caused a leftward shift in the E-S coupling curve (red vs. black symbols) that persisted upon PTX washout if HFS was performed in the presence of PTX (open symbols). D2. PTX without HFS: Application of PTX caused a leftward shift in the E-S coupling curve that was reversed upon PTX washout if no HFS was performed. Data shown represent the mean ± SEM.

In the absence of PTX, both the dendritic EPSP (Figure 3.A1) and population spike (Figure 3.A2) potentiated with HFS as expected. The first and second I/O curves are both pre-HFS, and the corresponding E-S coupling curves for these time points are similar, indicating a stable relationship between the dendritic EPSP and population spike (Figure 3.C1). After induction, the persistent leftward shift in the third curve shows that E-S potentiation was stable for at least three hours post-HFS (Figure 3.C1, open symbols). For comparison, when no HFS was performed, and in the absence of PTX, the E-S relationship was stable over the entire five hour recording period, as shown by the three identical E-S coupling curves at all three time points (Figure 3.C2).

The application of PTX to baseline recordings had various effects on the dendritic EPSP and population spike (Figure 3B). Application of PTX caused a very small increase in the dendritic EPSP (16 ± 4%; paired t-Test, p < 0.01) that could then be further significantly potentiated with HFS (paired t-Test, p < 0.01) (Figure 3.B1). Upon PTX washout, LTP was not different than LTP in the No Drug control (No Drug: 59 ± 10%, n=9; PTX-WO: 56 ± 7%, n=10). The population spike however, showed a substantial increase after PTX application (81 ± 14%; paired t-Test, p < 0.01) that could not be further significantly potentiated with HFS (Figure 3.B2). Upon PTX washout, the population spike remained potentiated at a level similar to that seen before HFS, and also at a level similar to that seen in the potentiated No Drug control (No Drug: 121 ± 7%; PTX-WO: 122 ± 13%). The resulting effect on E-S coupling from the application of PTX only can be seen by the leftward shift in the E-S coupling curve in the presence of PTX, but before HFS (Figure 3.D1, red vs. black symbols), and the emergence and persistence of E-S potentiation by the sustained leftward shift in the E-S coupling curve when PTX is washed out after HFS (Figure 3.D1, open symbols). In experiments in which no HFS was performed, and only PTX was applied, the observed increases in the dendritic EPSP and population spike could both be reversed to pre-drug levels upon PTX washout (EPSP: −6 ± 6%; PS: 6 ± 12%, n=9), and the resulting shift in the E-S coupling curve from PTX application (Figure 3.D2, red symbols) could also be reversed to a near pre-drug state upon PTX washout (Figure 3.D2, open symbols).

These results demonstrate that PTX mimics the expression of, but does not induce, E-S potentiation, and furthermore, that GABA_A_ blockade by PTX does not prevent the induction E-S potentiation by HFS. E-S potentiation can still be induced by HFS even in the presence of PTX, even though it is not visible in the presence of PTX. Thus, taken together, the experiments in Figures 1-3 show that blockade of GABA_A_ transmission does not block, but merely occludes, E-S potentiation.

The two most likely places where coupling between the dendritic EPSP and the somatic action potential can change are in the dendrite itself, between the synapses and the soma, and between the soma and spike trigger zone at the axon hillock, where synaptic inhibition may be the most influential. To isolate the electrotonic coupling between dendrite and soma, we compared measurements of the dendritic EPSP field with measurements of the EPSP field that appears at the soma, i.e., the initial rising slope immediately prior to the downward deflection of the population spike (see Figure 1A). This somatic EPSP represents the passive current source that is secondary to the active current sink driven by the dendritic synapses. It represents the amount of EPSP current that survives transit from the dendritic origin to the soma, and provides a way to measure dendro-somatic electrical coupling (D-S coupling), and has been used previously by others for similar analyses [5, 6, 15, 22, 23].

To determine if a change in D-S coupling does occur, and what the potential role of that change might play in the expression of E-S potentiation, we compared LTP of the dendritic EPSP field with that of the somatic EPSP field in the same conditions previously discussed: No Drug, PTX throughout, and PTX post-HFS (Figure 4).

**Figure 4:**
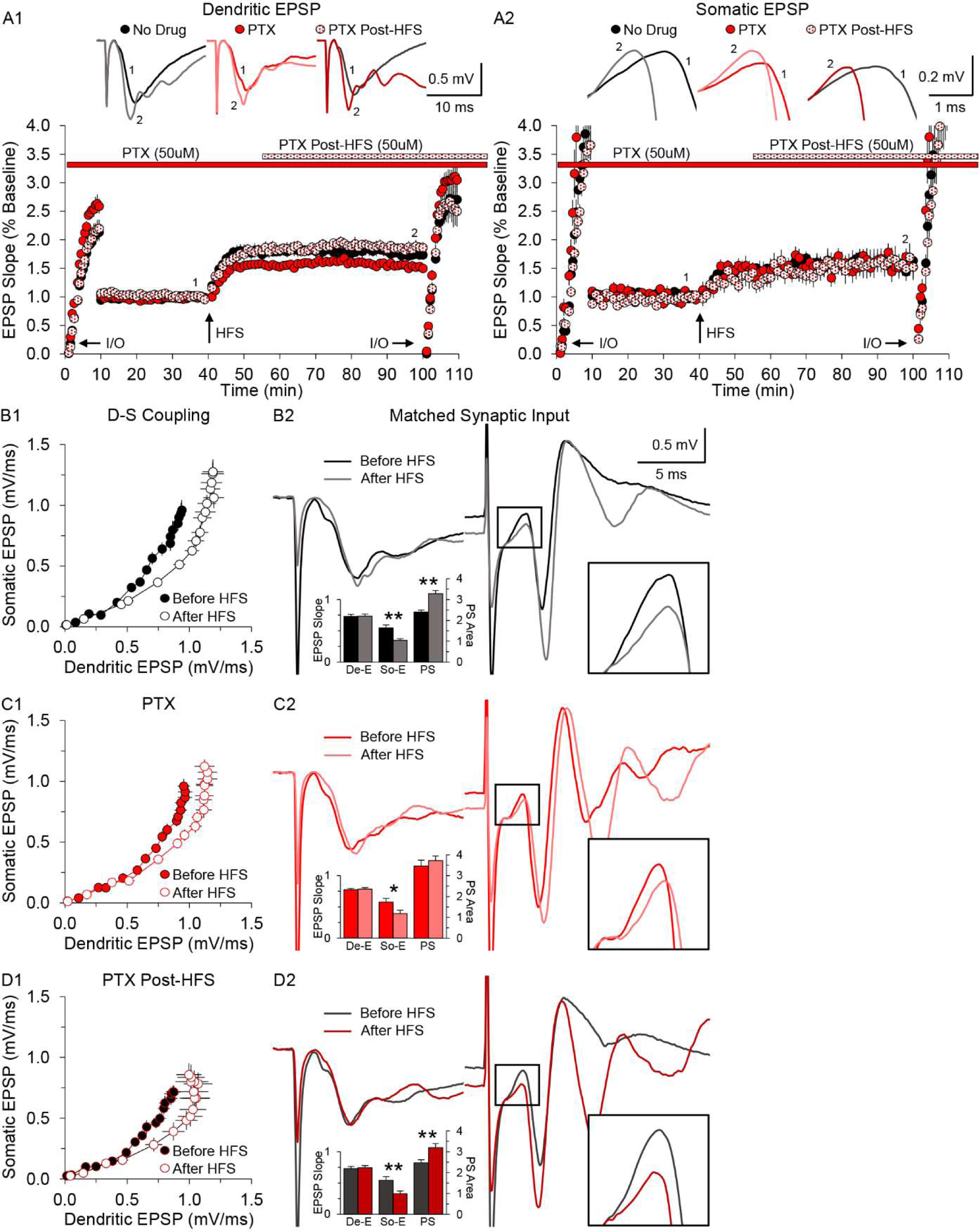
Dendro-Somatic (D-S) coupling is decreased after HFS. A. The time-course of experiments showing the effects of HFS (up arrow) on the dendritic EPSP field (A1) or somatic EPSP field (A2) in the presence of PTX (50 µM) applied either before or after HFS. Filled black symbols: No Drug control; red filled symbols: PTX present throughout experiment; red hatched symbols: PTX applied post-HFS. Time of delivery of PTX for each condition is indicated by the red bar (PTX present throughout experiment) or by the hatched bar (PTX applied 15 min post-HFS) located above the plot. An Input/Output curve was constructed at the beginning and end of each experiment (indicated by I/O arrows at the bottom of the plot) and used to construct the D-S coupling curves shown in parts B-D below. A1. Time-course of experiments showing the effects of HFS and PTX application on the dendritic EPSP field. Experiments are the same as in Figure 1.B1, and are shown here for comparison. A2. Time-course of experiments showing the effects of HFS and PTX application on the somatic EPSP field. The somatic EPSP field is the reflective field of the dendritic EPSP field, and is measured at the soma. Somatic EPSP slope data is taken from the simultaneous population spike field recording (see Figure 1A for reference). Long-Term potentiation of the somatic EPSP field was not different in any of the three conditions (No Drug: 56 ± 8%, n=12; PTX: 56 ± 14%, n=12; PTX post-HFS: 57 ± 21%, n=10). Averaged traces before (1) and after (2) HFS are displayed above the plots for each of the three conditions, and the corresponding time points for these averaged traces are indicated on the LTP time-course below. B1, C1, and D1. D-S coupling curves constructed by plotting the somatic EPSP slope against the dendritic EPSP slope (see Figure 1A for reference), data taken from the absolute values of I/O curves generated in the experiments shown above (also shown in Figure 1B). In all conditions there is a persistent shift in the D-S coupling curve to the right, not the left as seen with the E-S coupling curves. This indicates the electrical coupling between the dendritic input and the soma is decreased after HFS, opposite of what is observed with E-S coupling. B2, C2, and D2. Averaged traces of input matched dendritic EPSPs (left traces) and the resulting simultaneously-recorded population spikes with the somatic EPSPs highlighted and magnified in the boxed region (right traces) before and after HFS. Traces after HFS have a reduced stimulus strength to match the rising slope of the dendritic EPSP to that of the dendritic EPSP before HFS. In all conditions, matched dendritic EPSPs after HFS give rise to smaller somatic EPSPs compared to matched dendritic EPSPs before HFS. This results in the persistent rightward shift in the D-S coupling curve throughout the range of synaptic input as shown in B1, C1, and D1. The inset bar graphs show the show the significant reduction in the somatic EPSP (So-E) with matched dendritic EPSPs (De-E) after HFS compared to before (t-Test; No Drug and PTX post-HFS, p < 0.01; PTX, p < 0.05). The accompanying population spike data from Figure 2 is also shown here for comparison. Despite there being a reduction in the somatic EPSP for a matched dendritic input after HFS, there is still a larger population spike than before HFS with matched synaptic inputs in the No Drug control and PTX post-HFS groups. This is not seen when PTX is applied throughout. However, note that PTX does not influence the relationship between the dendritic and somatic EPSP fields as there is still a reduction in the somatic EPSP after HFS, suggesting that changes in GABA_A_ transmission has no, or minimal, participation in changes to D-S coupling. Data shown represent the mean ± SEM. Bar graph data are generated from matched dendritic EPSP slopes and resulting somatic EPSP slopes taken from Input/Output curves before and after HFS. Recorded traces for each matched pair were averaged and shown here in B2, C2, and D2 respectively for each group.

Potentiation of the dendritic EPSP shown here (Figure 4.A1) is the same as that shown in Figure 1.B1, and displayed again here for comparison. The somatic EPSP in all three conditions showed potentiation in response to HFS that was not different across conditions, but was overall proportionally less than potentiation of the dendritic EPSP.

D-S coupling curves for each of these three conditions were constructed as in earlier figures, by plotting the slope of the dendritic EPSP field potential against the slope of the somatic EPSP field potential across the full range of tested stimulus strengths. This shows the relationship between the dendritic synaptic input current and the amount of that current that reaches the soma, across a range of synaptic responses (Figure 4B-4D). The induction of LTP did, in fact, change the electrical coupling between dendrite and soma, but to our surprise, not in the expected direction. Instead of an increase in coupling, which would have been represented by a leftward shift in the coupling curve, we observed a rightward shift, indicating that the electrical coupling between dendrite and soma was reduced after the induction of LTP. This reduction occurred in the presence or absence of PTX, regardless of when it was applied (Figure 4.B1, C1, and D1). The averaged input matched traces shown here in Figure 4.B2, C2, and D2 are the same as those shown in Figure 2, with the somatic EPSP region boxed and magnified, and the addition of the respective somatic EPSP slopes from these traces quantified in the inset bar graphs below. The input matched dendritic EPSP traces clearly show a reduced resultant somatic EPSP after HFS in all three conditions, indicating a reduction in D-S coupling. This reduction is significant in all three conditions, even when PTX was present throughout and E-S potentiation was occluded (t-Test; No Drug: p < 0.01; PTX: p < 0.05, PTX post-HFS: p < 0.01). If PTX was absent during HFS, there was still a significant potentiation of the population spike afterwards, despite there being significantly less surviving current to the soma (Figure 4.B2 and D2). The averaged input matched traces shown in Figures 2, 4, and 5 represent a range of responses that demonstrate both the expression of E-S potentiation and the reduction in D-S coupling. Thus, our initial hypothesis that there would be an increase in electrotonic coupling was incorrect. This result shows there is a GABA_A_-independent influence in the dendrites of pyramidal neurons that is acting in opposition to E-S potentiation, in that the somatic EPSP is increased by LTP less than might be expected, due to the reduction in Dendro-Somatic coupling.

One possible mediator for this unexpected change in electrical coupling between dendrite and soma might be related to another form of synaptic inhibition, GABA_B_ receptor-mediated activity. GABA_A_ receptor density is greatest in the peri-somatic region, while GABA_B_ receptor density is more uniform, but greater in distal dendrites, and highest in the s. radiatum, where Schaffer collateral synapses terminate on CA1 pyramidal neurons [16, 24, 25]. To test the hypothesis that GABA_B_-mediated transmission was involved in the observed change in D-S coupling after HFS, we examined LTP, E-S coupling, and D-S coupling in the presence of the GABAB receptor antagonist CGP-54626 (CGP). Experiments were performed that were identical to those carried out using PTX as shown in Figures 1, 2, and 4, with the use of CGP (3 µM) in place of PTX (No Drug, n=12; CGP throughout, n=10; CGP post-HFS, n=10) (Figure 5). The No Drug control data used for comparison with CGP experiments is the same as that used with PTX experiments. Application of CGP did not have any detectable effect on the baseline coupling between the dendritic and somatic EPSP fields (D-S coupling), or dendritic EPSP and population spike fields (E-S coupling). LTP of the dendritic EPSP and population spike were not different across groups.

**Figure 5:**
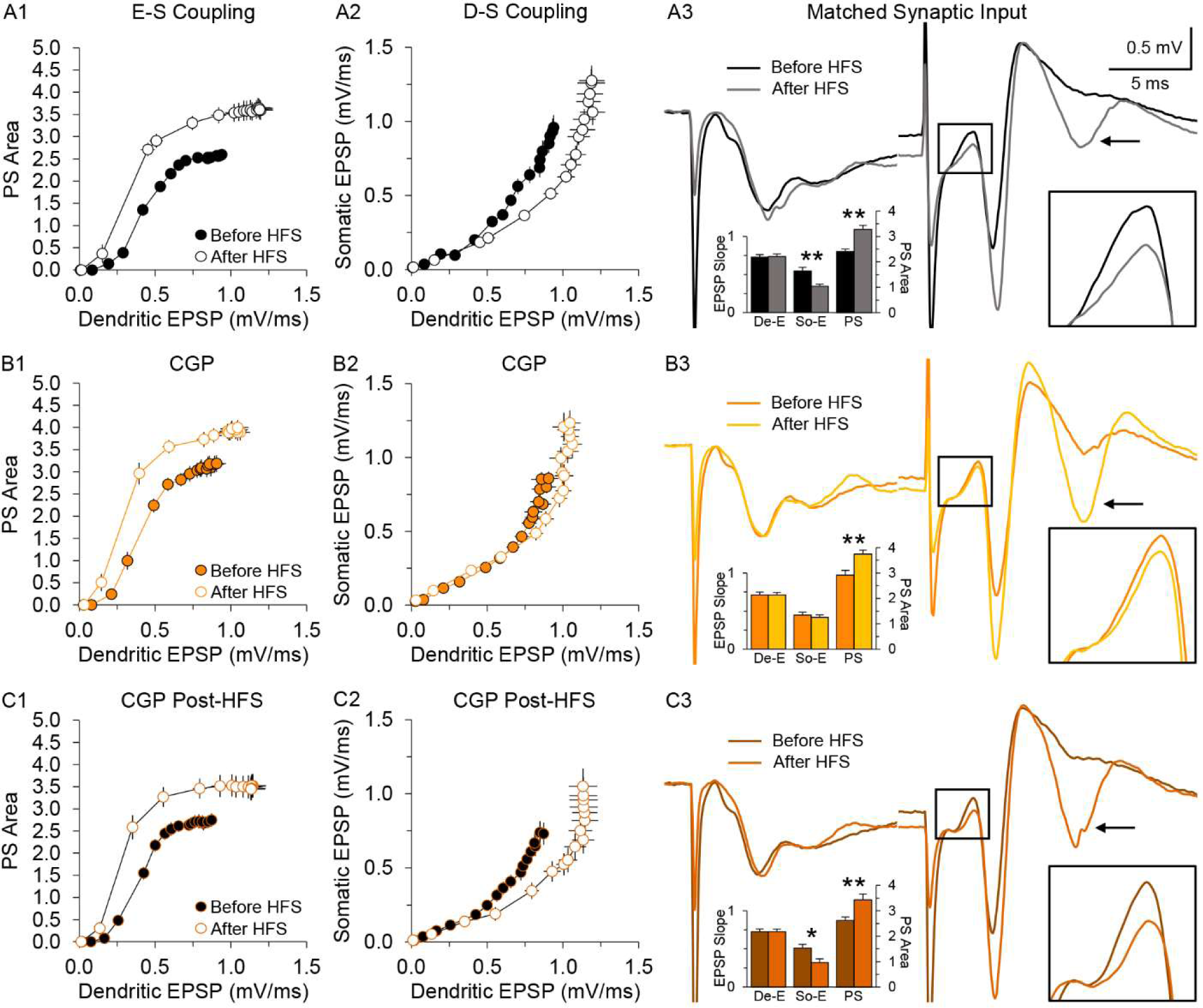
HFS-induced reduction in the induction, but not the maintenance or expression of Dendro-Somatic coupling, depends on the GABA_B_ receptor. Experiments were conducted identically to those carried out with PTX, with the GABA_B_ receptor antagonist CGP-54626 (CGP) (3 µM) used in place of PTX. All E-S coupling curves in this figure were constructed in an identical manner as those in the earlier PTX experiments. A. The No Drug control data (n=12) is the same as that shown in Figures 2 and 4, shown here for comparison. HFS resulted in a leftward shift in the E-S coupling curve (A1), and a rightward shift in the D-S coupling curve (A2). Matched dendritic EPSPs before and after HFS resulted in a smaller somatic EPSP (t-Test, p < 0.01) and larger population spike (t-Test, p < 0.01) after HFS compared to before HFS (A3), demonstrating EPSP-Spike potentiation and a change in D-S coupling. B1 and B2. When CGP was present during HFS, there was a persistent leftward shift in the E-S coupling curve (B1), but no change in the D-S coupling curve (B2), abolishing the rightward shift as was seen in the control. B3. Matched dendritic EPSPs give rise to somatic EPSPs that are not different, but generate larger population spikes, after HFS compared to before. This result agrees with the persistent leftward shift seen in the E-S coupling curve, but no shift in the D-S coupling curve, throughout the range of synaptic input (B1, B2). The inset bar graph shows no difference in the somatic EPSP (So-E) with matched dendritic EPSPs (De-E) after HFS (light orange bars) compared to before (orange bars)(t-Test, p = 0.55, n=10), which seems to have no effect on E-S coupling since the population spike (PS) was significantly potentiated (t-Test, p < 0.01) after HFS, indicative of E-S potentiation. C1 and C2. When CGP was applied after HFS, it had no effect on either E-S coupling (C1) or D-S coupling (C2), producing results similar to control. C3. Matched dendritic EPSPs before (brown bars) and after (dark orange bars) HFS gave rise to a smaller somatic EPSP (t-Test, p < 0.05, n=10) and larger population spike (t-Test, p < 0.01) after HFS, demonstrating EPSP-Spike potentiation and a change in D-S coupling similar to that seen in control. This suggests that GABA_B_ activity is required for the induction of the decrease in D-S coupling, but does not participate in its maintenance or expression. Note the afterdischarge spike(s), indicated by the arrows, in the population spike trace after HFS in all groups, thought to occur as a result of reduced inhibition following HFS (No Drug), or a combination of HFS and drug application (CGP, CGP post-HFS). Unlike the multiple afterdischarge spiking seen in PTX experiments, CGP seems to only enhance (visual observation, not quantified) a single afterdischarge spike. Data shown represent the mean ± SEM. Bar graph data are generated from matched dendritic EPSP slopes and resulting somatic EPSP slopes and population spike areas taken from Input/Output curves before and after HFS (Data not shown). Recorded traces for each matched pair were averaged and shown here in A3, B3, and C3 respectively for each group.

The effects of HFS on E-S coupling and D-S coupling in the presence of CGP were very different than that seen with PTX (Figure 5). When CGP was present during the induction of potentiation (HFS), the rightward shift in the D-S coupling curve was abolished (Figure 5.B2), but the leftward shift in the E-S coupling curve persisted (Figure 5.B1). The input matched traces when CGP was present throughout the experiment reflect these results (Figure 5.B3). One idea that may explain the rightward shift in D-S coupling observed in the control might be that GABA_B_-mediated inhibitory synaptic transmission is persistently enhanced after LTP induction, however when CGP was applied after induction, it did not reverse the rightward shift (Figure 5.C2), and both the E-S and D-S coupling curves remained shifted similar to that of the control (Figure 5A vs 5C). The input matched traces visually confirm that after HFS, there is in fact less surviving somatic EPSP (t-Test, p < 0.05), which still results in a larger population spike (t-Test, p < 0.01), similar to what is observed in control (Figure 5.A3 vs 5.C3). This result demonstrates that GABA_B_-mediated inhibitory transmission plays an important role in the decrease in D-S coupling following HFS, but that this role is restricted to the induction of this decrease, not in its maintenance and/or expression.

## Discussion

In this paper, we examined mechanisms underlying EPSP-Spike potentiation, the persistent increase in EPSP-action potential coupling that occurs alongside the more familiar Long-Term Potentiation (LTP). There are three obvious potential mechanisms that might support E-S potentiation: 1) A lowering of the action potential threshold at the postsynaptic spike trigger zone, 2) A greater survival of synaptic current as it moves from its dendritic origin to the spike trigger zone, or 3) A decrease in inhibitory synaptic shunts that usually serve to prevent current flow to the spike trigger zone. Here, we examine the hypothesis that E-S potentiation is a result of both a suppression of GABAergic transmission, and of potentially cell-autonomous changes in electrotonic coupling between dendrite and soma allowing for more efficient current flow from synapse to spike trigger zone.

Consistent with earlier findings, we found that application of the GABA_A_ open channel blocker Picrotoxin caused a maximal leftward shift in the E-S coupling curve, and there was no further leftward shift when HFS was subsequently applied [9]. This occurs because the area of the population spike, the measure of how many pyramidal neurons have been brought to threshold, is saturated by the application of PTX. However, we also found that this saturated leftward shift became persistent, not reversed, upon washout of PTX from the slice, only if HFS was applied. Together these findings show that PTX occludes the expression of E-S potentiation by maximizing the population spike, but does not actually induce E-S potentiation or prevent it’s induction by high-frequency synaptic activity.

While it is clear that GABA_A_ transmission participates in E-S potentiation in some way, and most likely represents one of the major mechanisms responsible for E-S potentiation, it is not clear that a persistent reduction of GABA_A_ transmission can account for all of its expression. Previous studies have suggested either a decrease in GABAergic signaling [6, 12, 14, 26, 27], no change in GABAergic signaling [27, 28], partial participation of GABAergic signaling (40%) [6], or even an increase in GABAergic signaling [4, 27, 29], many of these results in response to HFS. Nonetheless, in our own recordings we have multiple lines of evidence that suggest a suppression of inhibition specifically as a result of HFS as a major contributor to the expression of E-S potentiation, including the presence of multiple population spikes that are produced only after the induction of E-S potentiation, and are persistent for the duration of our experiments (see population spike traces, Figures 1-5). Multiple studies that have suppressed inhibition through a variety of pharmacological means, including our own results using PTX (Figures 1.B2, 2.B2 3.C2), demonstrate the emergence of after-discharge spiking as a direct result of suppressed inhibition [30–34].

The other major mechanism that might account for E-S potentiation is a change in efficiency of charge transfer from dendrite to soma. We measured this D-S coupling by comparing the dendritic EPSP field with the somatic EPSP field. Surprisingly, we found that tetanic stimulation was followed by a persistent decrease in D-S coupling, and that this decrease was prevented by application of the GABA_B_ receptor antagonist CGP-54626 when present during HFS. Thus, our original hypothesis that there would be an increase in D-S coupling was incorrect. We revised this hypothesis to suggest that this decrease in coupling was mediated by a persistent potentiation of GABA_B_ transmission to the dendrite, but we discarded this hypothesis when we showed that CGP application after HFS had no effect on the D-S coupling shift (CGP did not reverse the shift once it occurred). Thus, we now hypothesize that the decrease in dendritic coupling following tetanic stimulation is induced by GABA_B_ transmission, but is not maintained by it.

It is unclear presently how GABA_B_ transmission during the induction of LTP and E-S potentiation may be driving the observed change in D-S coupling, but the most likely explanation may involve changes in resting K^+^ conductance that may lead to greater shunting of electrotonic current from the synaptic origin to the soma. It is well known that GABA_B_ transmission is linked to K^+^ conductance, and shows both increases and decreases in conductance. GABA_B_ has been shown to be strongly linked to Kir3 activity [24, 35], which may be playing a role here. The other candidate that may explain our findings is a change in the cationic HCN channels that contribute to the I_h_ current. This channel has been suggested to change expression bidirectionally along the dendrites in response to various LTP induction methods [7, 8], and may shunt excitability in response to LTP induction.

In summary, we have shown that High Frequency Stimulation results in synaptic LTP, EPSP-Spike potentiation, and a reduction in Dendro-Somatic coupling. E-S potentiation is not visible in the presence of the GABA_A_ channel blocker Picrotoxin, but only because of pre-saturation of excitability prior to tetanic stimulation. The underlying mechanism for the induction of E-S potentiation however, can still be induced in this state, as its persistence is induced by HFS, even when additional potentiation of the population spike is occluded. That the separate GABA_B_-induced reduction in D-S coupling still occurs even in the presence of PTX, suggests that a suppression of GABA_A_ inhibition in the soma, but not the dendrites, may contribute to the expression of E-S potentiation. A reduction in D-S coupling does not occur in the presence of the GABAB receptor antagonist CGP-54626, but E-S potentiation does occur. This also suggests the locus of E-S potentiation is somatic, and that this mechanism is more powerful than the GABA_B_-induced decrease in dendritic coupling.

Figure 6 shows a proposed model to summarize and explain the results presented in this paper. Each panel shows a schematic representation of current flow with the width of each blue arrow indicating the decay in the amount of current as it travels from the dendritic excitatory synapses towards the spike trigger zone. Pre-HFS control conditions are on the left of each panel, and post HFS potentiated conditions are on the right, separated by the dashed line. In this model, current is shunted out through passive dendritic conductance (blue split-out arrows) and via synaptic inhibition (red arrows). GABA_B_ transmission may provide some neutralization/shunting in the dendrite of the excitatory synaptic current, but this does not change after potentiation. However, activation of the GABA_B_ receptor during HFS does appear to induce a process that persistently changes the passive dendritic shunt (gold arrows). The current that survives to the soma is subject to being neutralized by GABA_A_-mediated transmission, which is decreased persistently after HFS, leading to greater excitatory current reaching the spike trigger zone. Panel A illustrates the proposed changes after HFS with potentiation of the synaptic input, while panel B shows the same process, but when the synaptic input is potentiated, but then matched to the pre-HFS level by turning down the stimulus. The schematic width of the blue arrows is designed to show how HFS can cause a decrease in dendritic coupling and still result in greater action potential discharge at the spike trigger zone due to an over-balanced reduction in GABA_A_ transmission at the soma.

**Figure 6:**
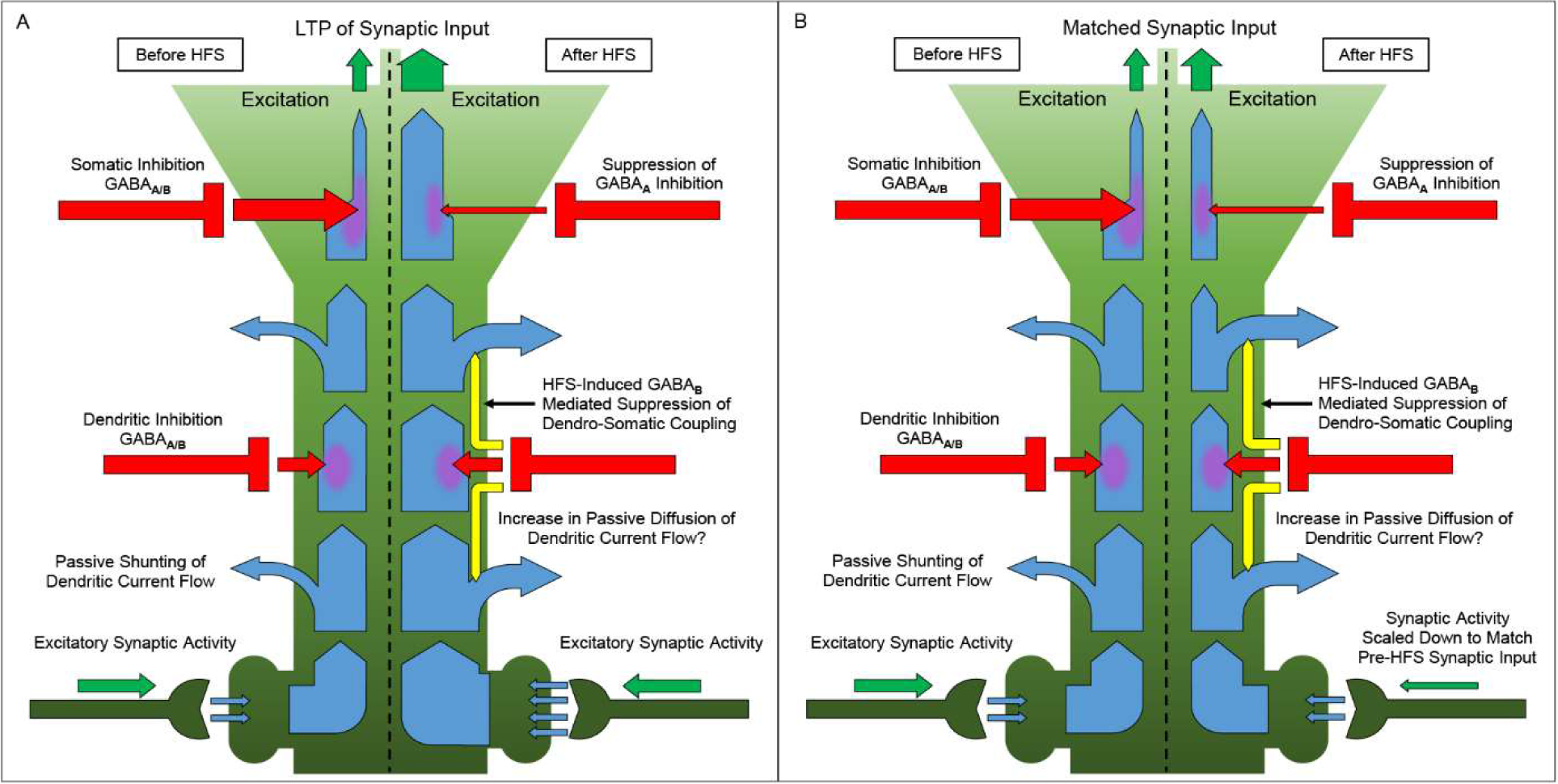
Changes in Dendro-Somatic electrical coupling accompany EPSP-Spike potentiation after HFS. Schematic illustrating the change in electrical coupling along the dendritic tree after HFS as excitatory current moves from synapses to soma, and the accompanying increase in excitability when this current reaches the spike trigger zone. Before HFS, excitatory synaptic current (blue arrows) is shunted along the dendritic tree as it moves from synapses to soma by passive dendritic current loss (blue arrows pointing outward), as well as blunting by GABAergic transmission (red arrows). The majority of inhibition however, occurs in and around the soma, and is largely driven by GABA_A_ transmission. GABA_B_ receptor activation during HFS produces an increase in electrical shunting through an unknown mechanism (yellow arrows) that persists after HFS, and is not dependent on GABA_B_ transmission for the expression and maintenance of this change. This occurs in conjunction with a suppression of GABA_A_ mediated inhibition in the soma, which allows for more current to reach the spike trigger zone, and results in an increase in excitability despite the reduction in D-S electrical coupling. A mechanism such as this may allow for regulating excitatory/inhibitory balance as a result of increased synaptic input and suppression of inhibition in the soma. A. Illustration of D-S coupling before and after HFS, where HFS produces an increase in excitatory synaptic transmission (LTP), along with an increase in electrical shunting as current moves up the dendrite to the soma. More current reaching the soma, in conjunction with a suppression of GABA_A_ mediated inhibition in the soma, results in greater excitability. B. Illustration of D-S coupling before and after HFS, where synaptic input after HFS has been scaled down to match synaptic input before HFS. After HFS, there is more electrical shunting, and less current reaching the soma compared to before HFS. However, due to a substantial suppression of GABA_A_ mediated inhibition in the soma, more current reaches the spike trigger zone, allowing for greater excitability compared to before HFS, and demonstrating the expression of EPSP-Spike potentiation.

A combination of LTP and E-S potentiation induced together, as illustrated in Figure 6, would serve to make the EPSP more effective in reaching action potential threshold. Since the final common path of a cell’s participation in neural information transfer is whether or not it emits an action potential, it is likely that the importance of E-S potentiation has been underestimated in the participation of plasticity of neural circuits, and its role in cognitive processes such as learning and memory. As a persistent increase in synaptic evocation of spike output, there is no reason to suppose that it could not participate in memory trace formation, and if it can be induced independently of LTP as has been suggested (2), then it could serve as an alternate path to such engram formation. Furthermore, if the decrease in dendritic coupling that accompanies E-S potentiation, previously unreported, could also be induced independently, something that has not yet been demonstrated, this could be a mechanism to weaken the expression of such engrammatic traces. In any case, it appears to be a mechanism by which a additional layer of the control of synapse-to-spike coupling can be modulated. These findings advance our understanding of information processing at a level of both the individual neuron, and the hippocampal microcircuit, and provide a better understanding and a greater appreciation of the role in E-S potentiation and the mechanisms that regulate its play in the information processing inside the neural circuitry.

## Acknowledgments

Funding for this work was provided by a grant from the National Institute of Mental Health (MH111768) and by the Harold and Leila Y. Mathers Charitable Foundation. We thank Sonja Winter from the Department of Psychological Sciences at UC Merced for helpful discussion about the use of statistics, and members of the Madison Lab for useful discussion on the experiments and manuscript.

